# Unstable Heteroresistance as a Stepping Stone to Stable Resistance via Compensatory Mutations

**DOI:** 10.64898/2026.10.08.756632

**Authors:** Brandon A. Berryhill, Dan I. Andersson

## Abstract

Antibiotic heteroresistance, a trait in which a drug-susceptible population harbors a rare resistant subpopulation, has been presented as an unstable intermediate state on the way to classic stable antibiotic resistance. However, the timescales and pathways under which this could occur has not been examined. Extending a recombination-based model of copy- heteroresistance to include compensatory mutations, with rates and costs from experimental estimates, we simulated populations across thousands of generations of growth with β-lactam exposure. In the first days a founder population survives by one of two paths, either amplifying the resistance gene to a high copy-number or gaining compensatory mutations and staying at low copy-number. However, both paths converge over weeks to months on the same low-copy compensated state, which would be indistinguishable from classical resistance. Importantly, the ending state would provide no information as to the route taken: an isolate sequenced afterward carries no sign of the intermediate amplified state. With the parameters estimated and across a wide range of parameters, we find that amplification-mediated heteroresistance can be a transient stepping stone across lethal selection that leaves no sequence trace in surviving isolates.

## Introduction

The choice of antibiotic to treat an infection essentially depends on one number, the minimum inhibitory concentration (MIC) of the infecting bacteria ^1^. That number is assumed to describe the whole bacterial population, but often it does not. In a phenotype called heteroresistance, a predominately drug-susceptible population carries a rare subpopulation which grows at higher antibiotic concentrations, often allowing them to survive antibiotic therapy ^2^. Heteroresistance is a common phenotype: with one report of 766 isolate-antibiotic combinations from 41 clinical isolates of four Gram-negative species tested against 28 antibiotics, more than a quarter were heteroresistant ^3^; and, another showing 88% of carbapenem-resistant Enterobacteriaceae to be heteroresistant to multiple antibiotics ^4^. However standard clinical susceptibility tests read only the dominant population, so they would call infections with these bacteria susceptible, with the resistant subpopulations undetected until the drug selects for them and treatment fails ^5–7^. Therefore, heteroresistance is a population-level phenomenon with the distribution of resistance states determining the dynamics of antibiotic treatment.

In Gram-negative pathogens, heteroresistance most often arises by tandem amplification of resistance genes encoding detoxifying enzymes ^8,9^. Repeated sequences such as insertion sequence elements flank the gene and recombination between sister chromatids during republication adds or removes copies of the gene. The number of gene copies a cell carries determines the amount of enzyme (e.g. β-lactamase) produced and thereby the MIC. Since the recombination event produces two daughter cells, one with increased gene copy number and the other with a decreased gene copy number, due to the fitness cost of gene carriage and expression, once the selective pressure is removed, the average copy number within the population can decline. While the per-copy fitness cost is relatively small, a cell carrying many copies of the gene replicates measurably slower than its low-copy-number kin. A heteroresistant population therefore can be understood as a community bearing a reversible trade-off where extra gene copies yield drug resistance at the cost of growth. The dynamics of the population under treatment is best captured by the behavior of the copy-number distribution. We have previously described a recombination-based model of this process which recapitulates the copy-number distribution of the subpopulations within a heteroresistant population and how they shift under selection—making quantitatively accurate predictions of the climb in copy number under drug selection and the partial reversion once the drug is removed ^10^. In that model, the only way to survive sustained drug treatment is to reach and hold high copy number at the population level by shifting the distribution of high-copy number states within the population. Therefore, a population that cannot increase average copy number fast enough is cleared by the treating drug.

Whether a population pays the full cost of that copy number climb, or finds a cheaper route, is presently unknown. As we know from numerous studies of traditional antibiotic-resistance, resistance does not have to stay costly ^11^. Across many resistance mechanisms, compensatory mutations, second-site changes that restore growth while keeping resistance, ameliorate that cost either partially or completely. Since heteroresistant cells are represented here as copy-number-variants, they have an route to compensation via a second-site mutation that confers resistance on its own at a potentially lower fitness cost. This second mutation would lower the number of copies a cell needs to survive and since there is a carriage-cost to these genes, average copy number should then decrease. A low-fitness-cost resistant state of this kind would compete with the costly high-copy climb beget by heteroresistance. This provides a potential mechanism to the adage that heteroresistance could be a transient state on the way to stable resistance ^12^. This argument has partial support: heteroresistant strains evolved under continuous drug selection begin to shed copies while staying resistant, so the descent starts, however the experiments generally stop there ^12^. They show a system moving toward a low-cost resistant state without showing which state it reaches, or how long arriving to this state takes. The endpoint of this descent decides whether heteroresistance converts fully to classical resistance and if the heteroresistance trait is lost from the population, and the timescale of that potential reversion has implications for antibiotic treatment of bacterial infections (and the design of experiments in the lab to mimic these therapies).

Here we ask whether heteroresistance is a stepping stone to stable resistance and investigate by what dynamics, on what timescale, and under what conditions this trait would serve as an intermediate state. To do so, we develop a mathematical and computer-simulation model extending our recombination-based model of copy-number heteroresistance to include compensatory mutations, with rates and costs from experimental estimates ^10^. We then follow heteroresistant populations from the first dose of a β-lactam across thousands of generations. We find that identical founding populations resolve stochastically into one of two outcomes at the hour and day timescales: i) average copy number climbs or ii) the population survives at low copy number by acquiring compensation mutants early. These outcomes occur in relatively fixed proportions across replicates. Interestingly, the high-copy state (outcome i) does not last. Selection drives copy number up within the first hours and days, but over the weeks and months that follow, average copy number is driven down due to the rise of compensatory mutants. Heteroresistance thus governs the early course of treatment and classical resistance governs the rest. The descent ends within the month timescale at a very-low-copy compensated state. Importantly, that state would be indistinguishable from classical resistance and an isolate sequenced at the end of treatment would carry no evidence of the heteroresistant amplification it climbed through. The compensation rate, the antibiotic dose, and the per-copy cost determine both the proportions of the two outcomes and the time to convergence of the states. Therefore, we conclude that heteroresistance is a transient route through lethal selection and which route a population takes is determined by a few measurable parameters.

## Results

### Mathematical and Computer-simulation Model of Recombination-based Heteroresistance

We developed a mathematical and computer-simulation model of heteroresistance under prolonged β-lactam treatment, extending our previous recombination-based heteroresistance model to include compensatory mutations ^10^. The model has twenty bacterial states: R0 through R9, the non-compensated states, which are the densities of heteroresistant bacteria carrying 0 through 9 copies of the antibiotic-resistance gene without a compensatory mutation, and R0C through R9C, their compensated counterparts. We elected to model 9 states as that is sufficient to cleanly differentiate low, medium, and high copy-number states while allowing for large state transitions (e.g., duplication from R4 to R8) without becoming too computationally intensive. Cells in either group recombine stochastically between copy-number states at the rates we experimentally estimated. The mechanism underlying this recombination are shown in Supplementary Figure 1. Each R*i* state can acquire a compensatory mutation at rate ω, transitioning to R*i*C, and revert to R*i* at the same rate. Treatment proceeds under a serial transfer regimen with a fixed antibiotic dose AMAX of 16 µg/mL delivered at each transfer after diluting all populations at a serial transfer rate of 0.01 (a 1:100 dilution). See the Supplementary Text and Equations, Supplementary Figure 2, and Supplementary Tables 1 and 2 for the details, a diagram, the states, and the parameters of the model.

We consider two scenarios for how the compensatory mutation affects bacterial susceptibility to the antibiotic: in the first, the compensatory mutation confers high-level resistance independent of copy number, and all compensated states share a single elevated MIC; in the second, the MIC of each compensated state scales with copy number, such that compensation and copy number determine resistance together (i.e., a copy-number-dependent MIC).

### Scenario 1: A single heteroresistant population resolves into one of two outcomes

Under a fixed β-lactam dose, a single heteroresistant founder reaches one of two outcomes. Following 4,000 replicate populations under Scenario 1, where compensation confers resistance independent of copy number, each resolved into one of these outcomes by 288 h, the span of a treatment course (Fig. 1A, B). In the first outcome, the population climbs to high copy number, where it survives albeit with a large cost to null fitness (Fig. 1A). In the second, the population stays at low copy number and survives by acquiring compensation mutations (Fig. 1B). Across all 4,000 replicate simulations, these outcomes occurred in roughly 85.5% and 14.5%, respectively (Fig. 1C), and each outcome was fixed well before the 288-h simulation horizon (Methods). No population went extinct at this dose. Extinction occurs only at very high antibiotic doses, where it becomes the dominant outcome (see the sensitive analysis).

**Figure 1.**
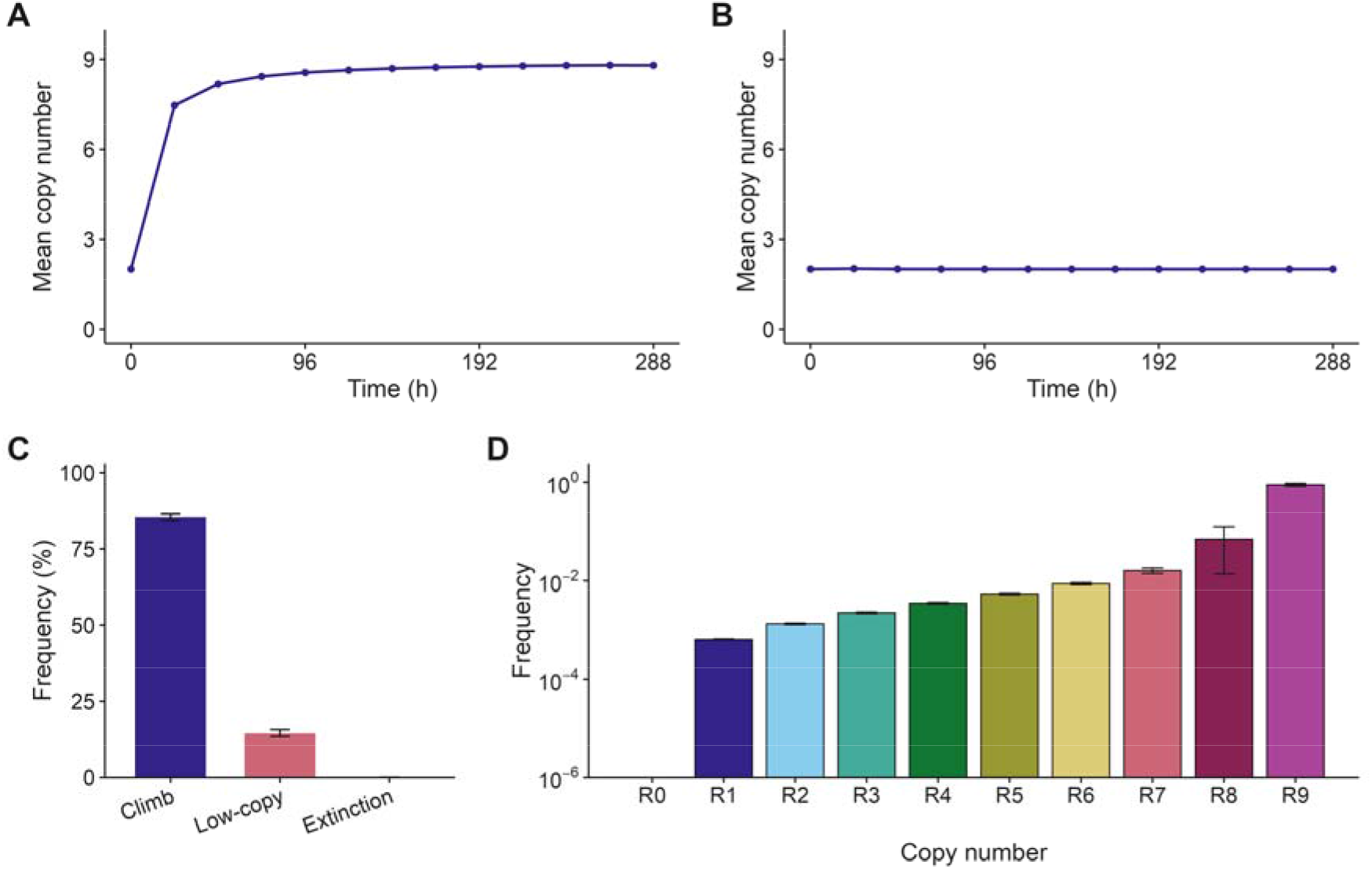
The outcomes of selection on a heteroresistant population under Scenario 1. *A)* Representative trajectory of a population that climbs to high copy number with a mean copy number near 8.8. ***B)*** Representative trajectory of a population that survives at low copy number through compensation with mean copy number near 2.0. ***C)*** Frequency of each outcome across 4,000 stochastic replicates, 85.5% climb, 14.5% low-copy compensated survival, and no extinction. Error bars are 95% Clopper-Pearson intervals. ***D)*** The ω = 0 control (no compensation): the copy-number distribution of the endpoint population at 288 h on a log frequency scale, shown as the mean and standard deviation across replicates.

The low-copy outcome requires the emergence of compensation mutants. With the compensation rate set to zero, every population climbed to high copy and the model reproduced the recombination-only model of its predecessor both in the endpoint distribution (Fig. 1D) and in the full dynamics (Supplementary Fig. 3). The base model already lets low-copy populations survive transiently through indirect resistance, in which high-copy cells degrade the drug. The low-copy-only survival outcome is therefore a product of compensation and not of indirect resistance.

The high-copy state does not last. Copy number rises fast, within the first 72 h, when the most amount of drug-mediated killing is occurring, until the heteroresistant nine-copy state R9 becomes dominant. However, compensated cells are present albeit at a low fraction (Fig. 2A). Compensation then sweeps through the high-copy population between roughly 350 and 500 h where the high-copy-number compensated state comes to dominate. Then selection begins to act on the relative fitness cost of carrying many copies of the resistance gene.

**Figure 2.**
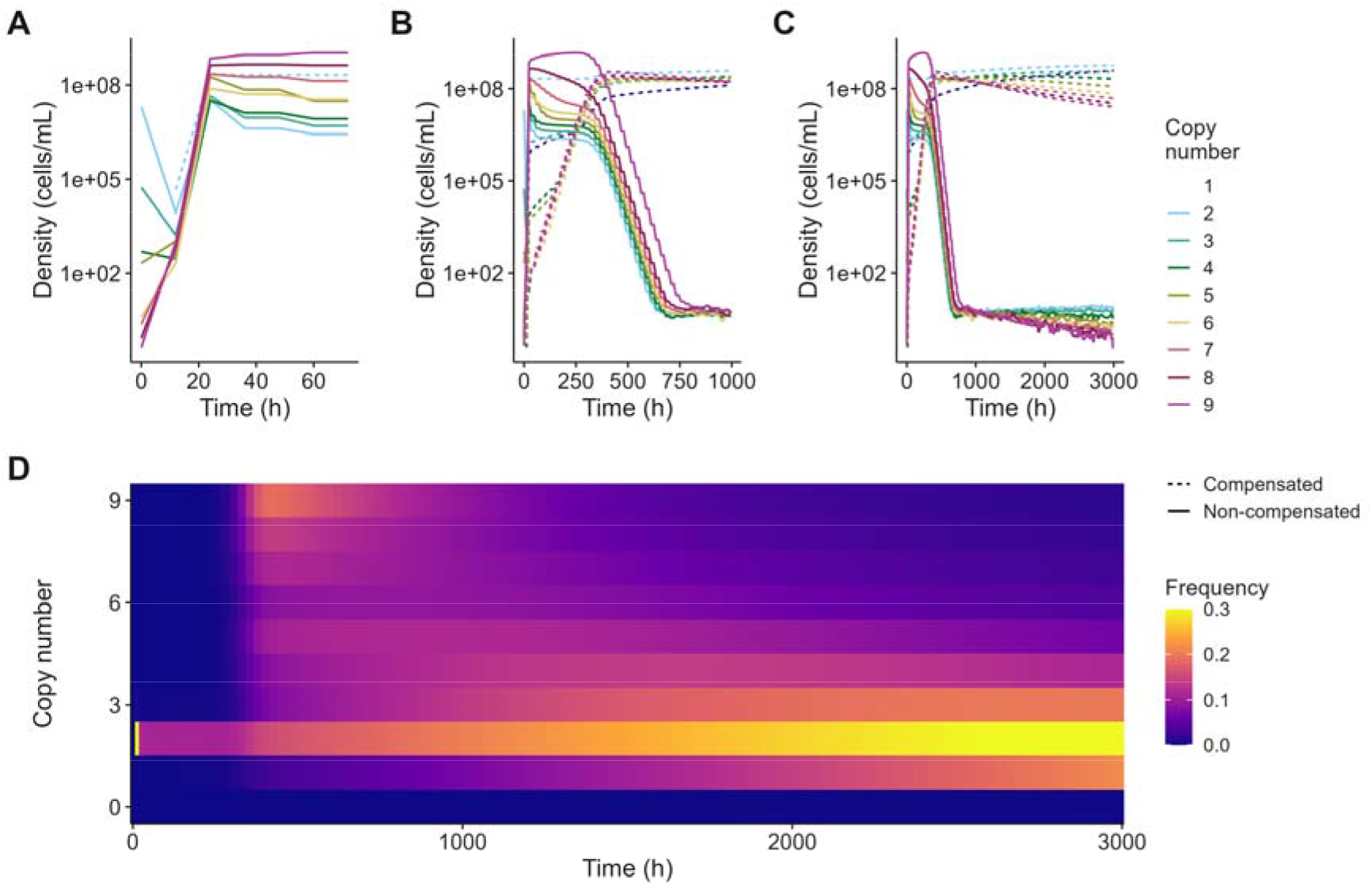
Simulations of the long-term dynamics of a heteroresistant population under Scenario 1 with one compensatory mutation. Copy number in A, B, and C are noted by color. Non-compensated states are solid lines and compensated states dashed. ***A)*** The heteroresistant climb over the first 72 hours, with compensation emerging. ***B)*** Dynamics over the first 1,000 hours, as intermediate-copy compensated cells come to dominate. ***C)*** Dynamics over 3,000 hours, as two-copy compensated cells take over. ***D)*** Heatmap of copy-number frequency over time, with the dominant copy number descending piecewise as the population shifts to lower copy.

The descent in average copy number occurs piecewise. Each copy lost lowers the fitness cost, so within the compensated mutants lower-copy cells outgrow higher-copy ones, and the recombination between sister chromatids supplies those lower-copy states for selection. By 1,000 h intermediate-copy compensated cells dominated (Fig. 2B), and by 3,000 h two-copy compensated cells have taken over (Fig. 2C), with the piecewise nature of this selection clearly visible (Fig. 2D). The descent did not finish within the 3,000h simulated window. The single-copy compensated state overtakes the rest only after about 4,500 h and the zero-copy compensated state R0C, the resistant endpoint that carries no amplification, overtakes single copy only beyond 126,000 h (Supplementary Fig. 4). The descent stops short of zero for most of the time course as losing the last copy occurs by a rarer recombination event than the steps between higher copies. The climb and the descent occupy different timescales, with the climb occurring over hours and days and the descent over the weeks and months that follow.

### Scenario 2: Copy-number-coupled compensation prevents descent

Coupling compensated resistance to copy number prevents the descent entirely. In Scenario 2, each compensated state’s MIC is three times the heteroresistant value at the same copy number. That protection gain exceeds the small per-copy cost at every step, so net growth under drug-exposure increases all the way up the copy number ladder such that selection favors the highest copy number rather than the lowest. As in Scenario 1, the population climbed and acquired compensation within the first hours, reaching the nine-copy compensated state by about 500 h (Fig. 3A), but from there it held at that copy ceiling for the rest of the run rather than descending (Fig. 3A, B).

**Figure 3.**
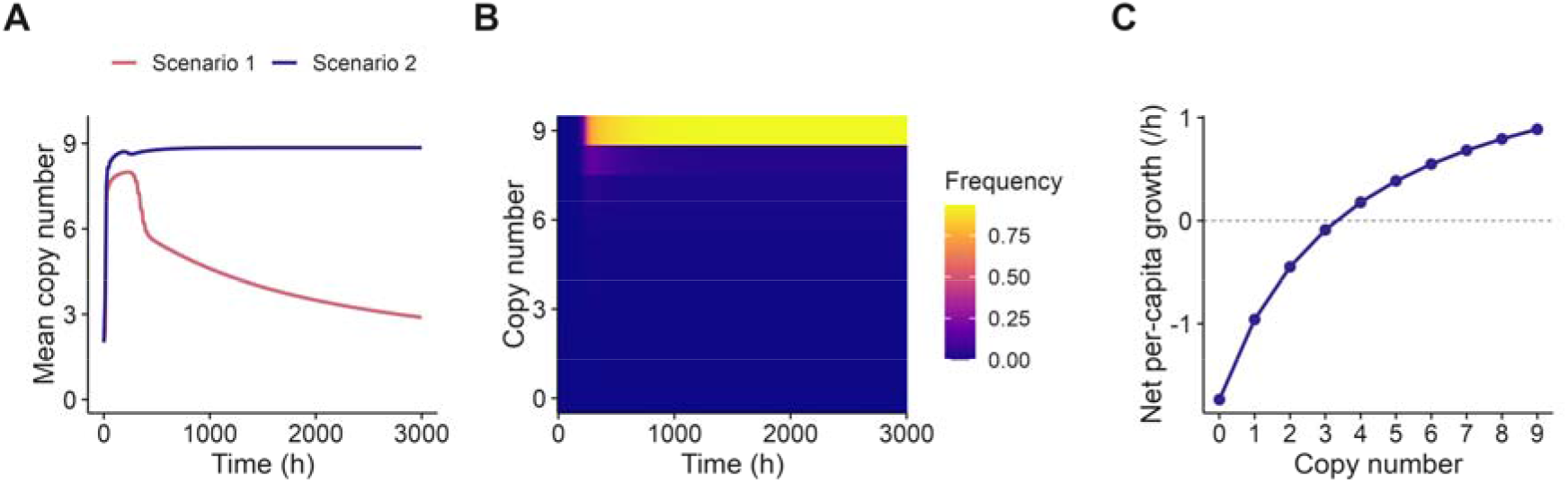
Scenario 2, copy-number-coupled compensation. ***A)*** Mean copy number over time for Scenario 1 and Scenario 2 overlaid. ***B)*** Heatmap of copy-number frequency under Scenario 2. ***C)*** Net per-capita growth as a function of copy number under Scenario 2.

No population descended and none were cleared, every replicate climbed and held at the copy ceiling, in 100% of replicates. As killing falls with each added copy, losing a copy lowers net growth even though the cell would still survive the dose, so selection favors the population at the highest copy number (Fig. 3C). Whether a population descends to low-copy resistance or holds at high copy therefore depends on whether compensation acts independently of copy number, as in Scenario 1, or scales with it, as here.

### Sensitivity Analysis: Antibiotic dose and compensation rate determine the outcome frequencies

Six parameters in the model determine how often each outcome occurs. To determine the extent to which the model is sensitive to these parameters, we varied the antibiotic dose, the compensation rate, the per-copy fitness cost, the recombination rate, the serial transfer rate, and the fraction of the cost that compensation recovers and measured the influence of each on the frequencies of the three outcomes (stable low copy number, a climb through high copy number, or clearance of the bacteria by the drug) using PAWN ^13–15^, a density-based measure of how strongly each parameter shifts the distribution of outcomes (Fig. 4, Methods).

**Figure 4.**
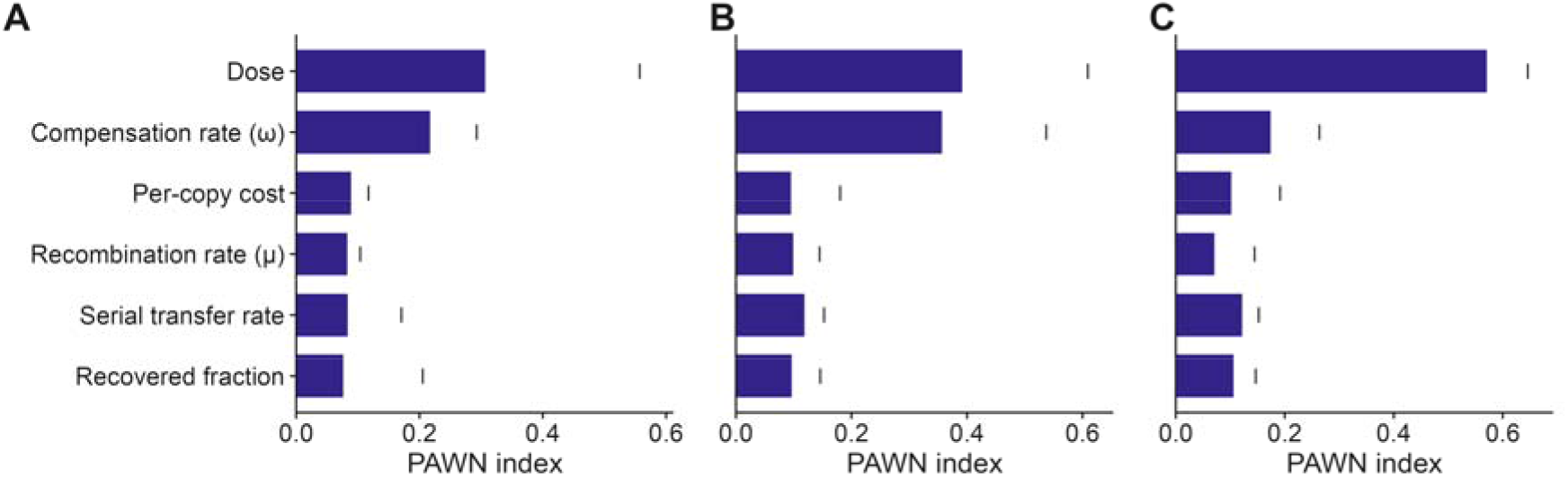
Global sensitivity of the three outcome frequencies to six model parameters: the antibiotic dose, the compensation rate, the per-copy fitness cost, the recombination rate, the serial transfer rate, and the fraction of cost recovered by compensation. PAWN sensitivity indices for each parameter against ***A)*** the climb-to-high-copy frequency, ***B)*** the low-copy compensated frequency, and ***C)*** the extinction frequency. Each bar is the median PAWN index for that parameter, and the tick marks the maximum index across the conditions.

The antibiotic dose was the strongest driver of all three outcomes, and it was the dominant control of the extinction outcome: clearance occurred almost entirely at the higher doses of antibiotics (Fig. 4C). The compensation rate was the next strongest driver of the two survival routes, the climb to high copy and low-copy compensated survival, and it had little effect on extinction (Fig. 4A, B). The remaining parameters, the per-copy fitness cost, the recombination rate, the serial transfer rate, and the fraction of cost recovered, each had only a small effect on any outcome. The per-copy cost, which determines how fast the descent occurs, did not measurably shape which outcome a population reached. The raw parameter-outcome relationships behind these indices are shown in Supplementary Fig. 5.

## Discussion

Heteroresistance can serve as a transient route to stable resistance, however if stable resistance progresses through heteroresistance or emerged directly can be genetically erased such that it is difficult to determine retrospectively. A founder population that survives sustained drug exposed climbs to high copy number, then walks back down to lower copy number as compensation mutants arrives, before ultimately settling into a very low-copy number compensated state indistinguishable from classical resistance. A low-copy compensated cell looks like any other resistant cell, so an isolate sequenced after treatment gives no sign of the heteroresistant amplification that occurred to reach the final state. A clinically important implication of this is that the heteroresistant history of a clinical isolate can only be recovered from sequencing data by longitudinal sampling of a treated patient. Experimental evolution has shown that populations lower their average copy number once compensation occurs ^12^, but these experiments have not followed these populations to a resting state. Our model predicts the final steady states, the time to reach it, and the frequency of populations ending up in those final states.

The two phases of survival through lethal selection occupy different timescales. The climb happens in the first hours and days of treatment, while drug killing is strongest which determines whether a population survives at all. The descent to single-copy resistance plays out over weeks, months, and longer, governed by the small growth advantage. The per-copy fitness cost determines the speed of the descent with a larger cost steepening the growth difference between adjacent copy states and shortening the time to reach the single-copy endpoint. Therefore, we can say that heteroresistance governs the early, decisive part of treatment and classical resistance determines everything after.

More than a quarter of pathogen and antibiotic combinations tested are found to be heteroresistant ^16^. However, if heteroresistance is instead a transient stage on the route to stable resistance, that frequency describes populations caught at different points along their evolution. Some pathogens scored susceptible may be early in this trajectory, carrying a resistant subpopulation that treatment will select and then convert into stable resistance over the course of therapy. Contrariwise, some pathogens determined resistant may have emerged from heteroresistant populations. The sole factor governing this disappearing-intermediate dynamic is the interaction between the compensatory mutations and the copy-number. Stated another way, the route a population takes is set by the kind of compensation mutants available to it.

The utility of theory comes from its ability to make testable predictions and the predictions presented here can be tested experimentally. In the laboratory, a serial-transfer evolution experiment that starts from a heteroresistant strain and tracks the copy-number distribution of the surviving population over thousands of generations would show whether the descent occurs, where it ends, and how long it takes. The clinical data which would support or reject the model requires the conditions under which the descent has time to run. For example, a heteroresistant infection which was initially scored susceptible by standard testing, in a site where immune clearance is weak such as the urinary tract, the prostate, or in an immunocompromised host, that is then treated with a long course of a β-lactam ^17–20^. In theory, this is the optimal setting in which a population could climb, compensate, and descend all within a single patient.

The model employed has four primarily limitations: First, its parameters come from only one system. Albeit the sensitivity analysis did not identify extreme sensitivity to any of the system-dependent parameters. Second, this model does not account for the impact of the innate immune system on treatment outcome ^21^. Innate immune effectors such as phagocytes could well control the emergence of rare resistant populations and prevent the ascent up the copy number ladder ^22^. Third, the serial transfer regimen at a fixed dose represents an idealized treatment which does not account for the pharmacokinetics of the treatment agent and the fluctuating drug concentration that would occur in vivo ^23^. Finally, the low-copy compensated endpoint is the resting state across the treatment-relevant timescale, but it is not the final steady state of the system. Far beyond any reasonable clinical horizon, the model predicts that the last copy will eventually be lost and a zero-copy compensated state will take over. Taken together, these limits do not lower the value of the model, nor do they drastically impact the predictions. We do not claim this model to quantitatively model the dynamics of treatment, but rather we claim the hypotheses generated from the theory are testable.

While our model and interests lie in antibiotic resistance, the principle described here is more generally applicable. Wherever a reversible amplification carries a population through a transient pressure and is then resolved either by compensation, copy loss, or divergence of the amplified copies, these dynamics could well occur. The same loci expansion and contraction appear in poxvirus adaptation to host defenses, where a gene accordion amplifies an anti-host factor and then contracts once an adaptive point mutation arises ^24^. The same is seen in extrachromosomal oncogene amplification in tumors, where copy number rises under targeted therapy and falls when the drug is withdrawn ^25–27^. Tumors under targeted therapy amplify the drug’s target or a bypass oncogene, and these focal amplifications are often unstable and can be lost once a stable resistance mechanism takes hold ^28^. In each case, amplification serves as a transient strategy to survive lethal selection which is quickly lost once a low-cost solution appears.

In conclusion, heteroresistance is best understood not as a stable trait, but as a transient route across lethal selection that can end in a resistance indistinguishable from any other and the benefits of this trait to survival are best seen in the short-term, with standard resistance governing the longer-term dynamics of antimicrobial treatment.

## Methods

### Numerical Solutions (Simulations)

We solved the model by fixed-step Euler integration of the coupled differential equations (Supplementary Equations 3 to 25), with a Monte Carlo transition applied at each step ^29,30^. The deterministic part of the equations controls the growth of the twenty population densities, the consumption of the limiting resource, the antibiotic concentration, and the β-lactamase production. The stochastic element governs the transitions between states, recombination between copy-number states at rates μ and γ, as well as compensation between each state and its compensated counterpart at rate ω, by drawing a random number against each transition’s rate at every step, as described in the Supplementary Text. Treatment proceeds under a serial transfer regimen where at each 24 h cycle all states and the antibiotic are diluted 1:100, the β-lactamase is reset to zero, the resource is restored to its maximum, and a fresh antibiotic dose is delivered. We followed replicate populations from identical initial conditions, differing only in their random draws, to the 288-h classification horizon. We ran longer stochastic batches to 3,000 h to follow the descent shown in Fig. 2 and Fig. 3, and a single long deterministic run to follow the far-horizon dynamics shown in Supplementary Fig. 4.

### Route Classification

We assigned each replicate to one of three outcomes at the horizon from two quantities. The first is the total population which is the sum of the densities across all twenty states. The second is the mean copy number, the population-weighted mean of copy number across the twenty states which is the sum of copy number times density divided by the total density. We score the total population size at the 288-h horizon, but we read the mean copy number as its peak by determining the average copy number at the each of the 12 transfers such that a population that climbed to high-copy and then began to decrease in average copy number within the first 288 h is still noted as having climbed to a high-copy number. A replicate whose total population at the horizon falls below one cell is labeled as extinction. If the replicate did not go extinct, then a peak copy number is scored. A simulation whose peak mean copy number reached five or more is scored as a climb to high copy, otherwise it is scored as low-copy compensated survival. The threshold of five copies lies between the two attracting states of the dynamics, near eight to nine copies for the high-copy outcome and near two for the low-copy outcome.

### Simulation engine and verification

We implemented the model in two forms. The first is an R model, written as a hand-coded fixed-step Euler loop with the per-step Monte Carlo transitions, extending the published recombination model to add the compensated states and the ω compensation transitions ^31^. It was built for validation of the second AI-generated model and for access to sensitivity-analysis packages in R. The second is a C++ model, which was used to generate the data presented here ^32^.

We verified the AI-generated C++ model against the R model in two ways, first at the level of trajectories and then on the outcome distributions. The first validation is completely deterministic. With the Monte Carlo generation disabled in both models the simulations are a deterministic Euler integrator of the model’s equations. We compared the two models’ outputs at within and between transfer densities across all twenty-five output columns: time, the twenty state densities, the resource, the antibiotic, the β-lactamase, and an index which counts the transfer number. The relative error was 1.76 • 10^-16^ at the first Euler step, 3.49 • 10^-14^ over a 24 h trajectory prior to transfer, and a maximum of 1.313 • 10^-13^ across all twenty-five output columns over a 288-h trajectory. However, this did require a compiler flag disabling fused multiply-add (-ffp-contract=off) to match the operation order of the C++ model to that of R.

The second validation focuses on the distributions with the stochastic transitions. With the Monte Carlo transitions enabled, the two implementations are not expected to agree cell for cell because they draw from different pseudorandom number generators, R’s uniform generator and the C++ model’s xoshiro256++. Therefore, we compared outcome frequencies instead by running the C++ model for 4,000 replicates and the R model reference for 100 (this corresponds to roughly the same compute time for each model). For each of the three outcomes we computed an exact 95% Clopper-Pearson binomial confidence interval found the C++ interval to overlap the R reference interval ^33^.

### Sensitivity Analysis

We measured the influence of six parameters on the outcome frequencies under Scenario 1 using the given-data variant of PAWN, a density-based global sensitivity analysis ^14,15^. PAWN scores an input by the distance between the unconditional output distribution and the distribution conditional on the input of interest, so it assumes only that the output is a distribution and not that variance captures it. The given-data estimator computes this from a single sample rather than a conditional sampling design by partitioning the output into bins by the quantiles of each input. The sensitivity index is therefore the distance between each bin’s output distribution and the unconditional distribution. The six inputs were the antibiotic dose AMAX, the compensation rate ω, the per-copy growth-rate cost, the recombination rate μ, the serial transfer rate, and the fraction of the cost that compensation recovers. We drew 300 parameter sets by Latin hypercube sampling with ω from 1•10^-10^ to 1•10^-6^, μ from 1•10^-7^ to 1•10^-3^, and the serial transfer rate from 1•10^-3^ to 1•10^-1^ each sampled on a log scale; AMAX from 4 to 64; the per-copy cost from 1•10^-4^ to 1•10^-2^; and, the compensated fraction from 0.1 to 0.9 about a baseline of 0.5. All other parameters were held at the values in Supplementary Table 2. For each parameter set we ran 100 stochastic replicate populations to the 288-h classification horizon and recorded the frequency of the three outcomes. For each input we partitioned the 300 sets into bins by the input’s quantiles and computed the Kolmogorov-Smirnov distance between each bin’s outcome-frequency distribution and the unconditional distribution, taking the median across bins as the sensitivity index (Fig. 4). Sampling used the lhs package in R, and the PAWN given-data indices were computed ^34,35^.

### Use of Artificial Intelligence

Claude Opus 4.8 (Anthropic) was used to convert the model from R to C++, which reduced the runtime of the model by roughly an order of magnitude ^36^. The converted code was verified against the R reference before any results were generated from it, as described above. The authors reviewed all model outputs using non-artificial intelligence.

## Acknowledgments

The authors would like to thank Matthew Lukenge, Sofia Jonsson, and Hervé Nicoloff for their role in estimating the parameters used in this study as well as Rotem Gross for his discussion of the results. Brandon would like to thank Amar Mujkic for convincing him to use AI to convert the R program to C++ rather than trying to do it by hand. This work was funded by grants to DIA from the Wallenberg Foundation (2018.0168), the Swedish Research Council (2021-02091), and the NIH (1U19AI158080-01). Open access funding was provided by Uppsala University.

## Data Availability

The data underlying all figures are provided as a Source Data file with this article and can be regenerated from the code below.

## Code Availability

All code used in this study, including the C++ simulation engine, the R reference implementation, and the analysis and sensitivity-analysis scripts, is archived on GitHub, but is private as of the time of this BioRxiv submission.

## Supplementary Materials

## Supplementary Text and Equations

### Pharmacodynamics of Antibiotic Treatment

We assume there is a single limiting resource (*r*, μg/mL) which in the absence of antibiotics would be the sole limiting factor to the growth and ultimate density of the bacteria. The pharmacodynamics of the antibiotic and treated bacterial populations is given by a Hill function where *Π(A,r)* is the net growth/death rate of the bacteria ^37^. Supplementary Equation 1 is written generally and applies to each of the twenty bacterial populations independently. To simulate the effect that the decreasing resource concentration has on the physiological state of the bacteria and their interactions, we include a term *ψ(r)* given in Supplementary Equation 2^38^.

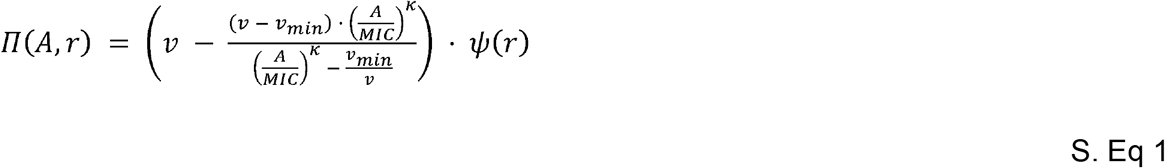

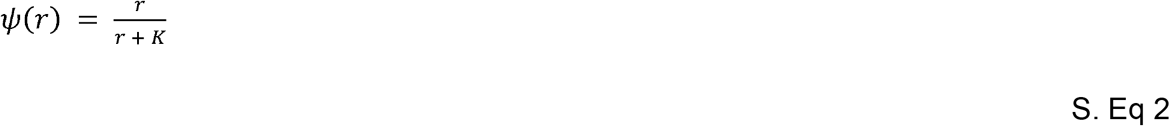

### Stochastic Transitions Between States

We simulate the transitions between states stochastically using a Monte Carlo process (18139350). A random number *x* (0 ≤ *x* ≤ 1) is generated from a rectangular distribution ^39^. If *x* is less than the product of the number of cells in the generating state (e.g., R3, CFU/mL), the volume of the vessel (Vol, mL), the transition rate (μ for the most frequent recombination events, γ for less frequent recombination events, or ω for compensatory mutations, all per generation per cell), and the step size (dt) of the Euler method employed for solving the differential equations then a transition occurs ^40^. For example, if *x < R3·Vol·μ·dt*, then MR3R2 cells are added to the R2 population and removed from the R3 population, where MR3R2 is 1/(dt·Vol). The same procedure is used for transitions between C states (e.g., MR3CR2C) and for compensation transitions between an R state and its C counterpart (e.g., MR3R3C and MR3CR3).

### Heteroresistance Model

This model is adapted from the model presented in ^10^. Briefly, the bacteria transition between twenty different states: R0 through R9, the densities (in cells per mL) of bacteria carrying 0 through 9 copies of β-lactamase gene, and R0C through R9C, the densities of their compensated counterparts which have additionally acquired a compensatory mutation that restores 50% of the fitness cost incurred by carrying the resistance gene. Transitions between R states occur stochastically as described above; transitions between C states occur via the same mechanism mirrored across the compensated space. The ability to transition between R states (or between C states) is limited by the copy number of the antibiotic-resistance gene present on each chromatid. The rate of transition is proportional to the number of ways recombination events can generate daughter cells with that copy number. Each state R*i* can additionally acquire a compensatory mutation at rate ω, transitioning to its compensated counterpart R*i*C, and revert at the same rate. A diagram of the model is presented in Supplementary Figure 2. The mechanism and rates of transition between R states (mirrored identically for the C space) are reprinted from ^10^ in Supplementary Figure 1. This model assumes serial transfer conditions whereby all bacterial states and the antibiotic are diluted 1:100 every 24 hours, the β-lactamase is reset to zero, the limiting resource is reset to its maximum, and a fresh dose of antibiotic at concentration AMAX is delivered. Given these assumptions, the pharmacodynamics above, the definitions of the states (Supplementary Table 1), and the parameters and their values (Supplementary Table 2), the rates of change in the densities of the different populations are given by Supplementary Equation 3 through Supplementary Equation 25. β-lactamase production in Supplementary Equation 5 is scaled by the conversion efficiency *e*, the same physiological factor that sets resource consumption in Supplementary Equation 3.

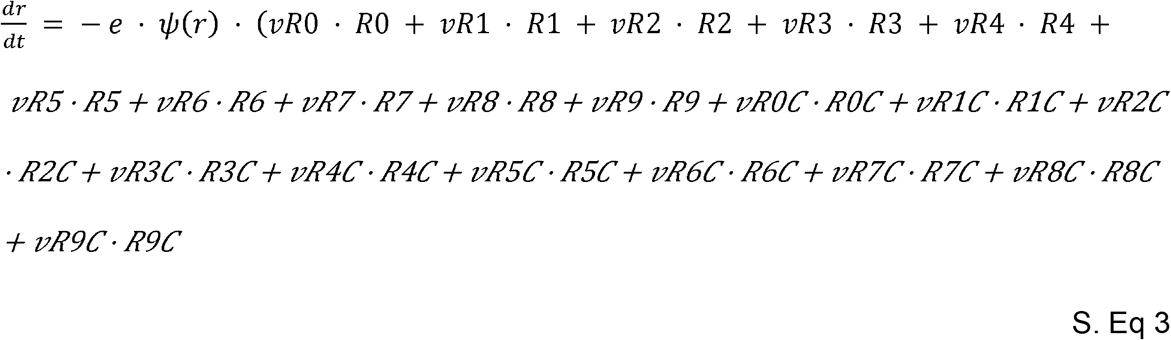

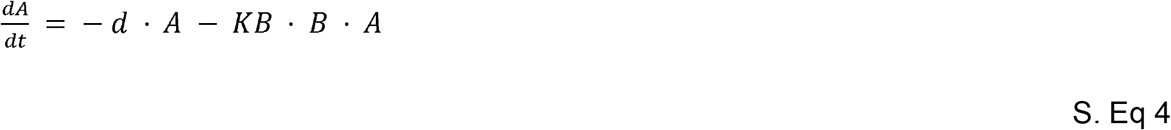

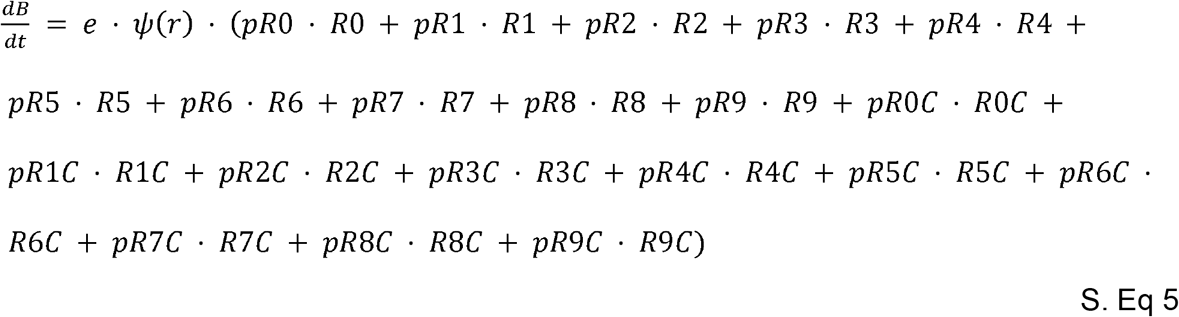

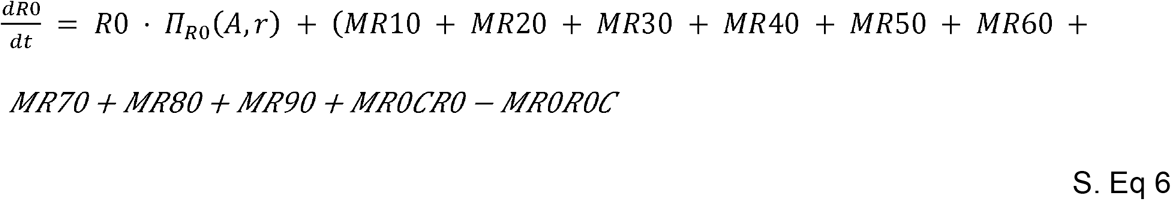

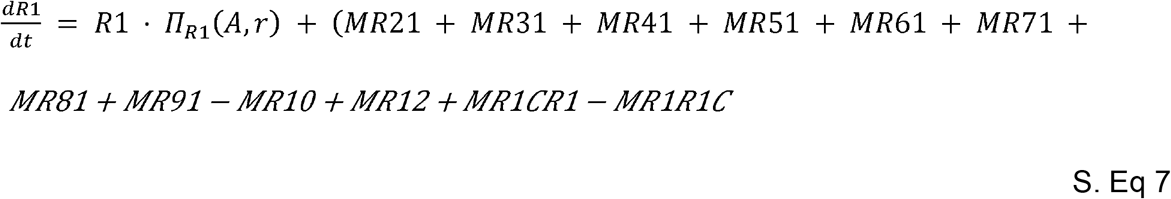

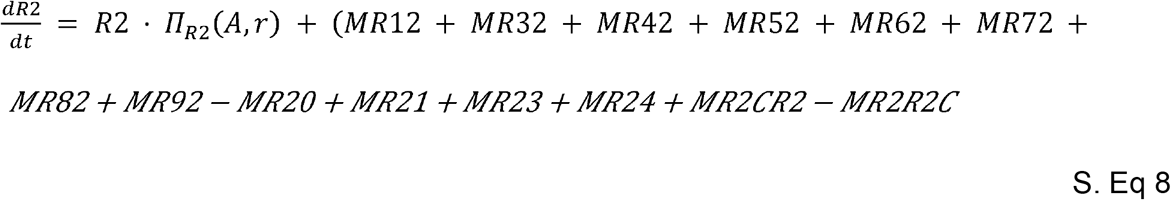

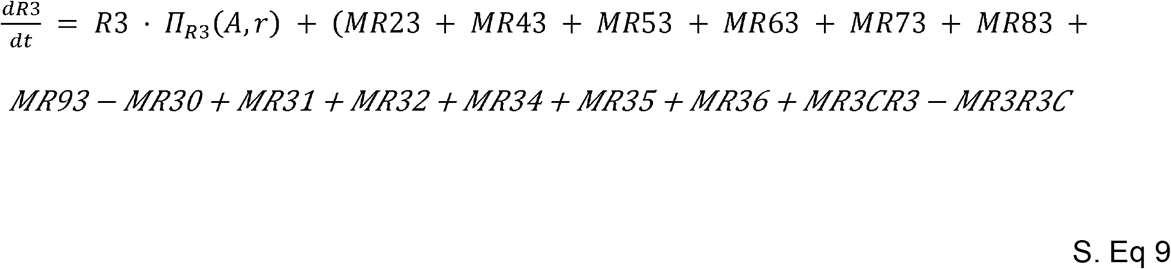

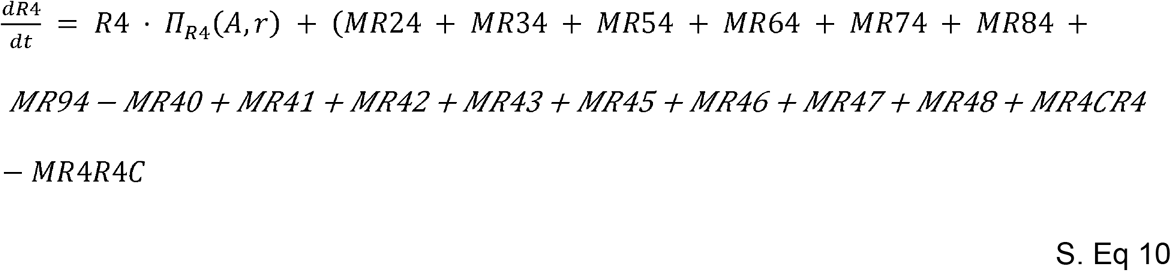

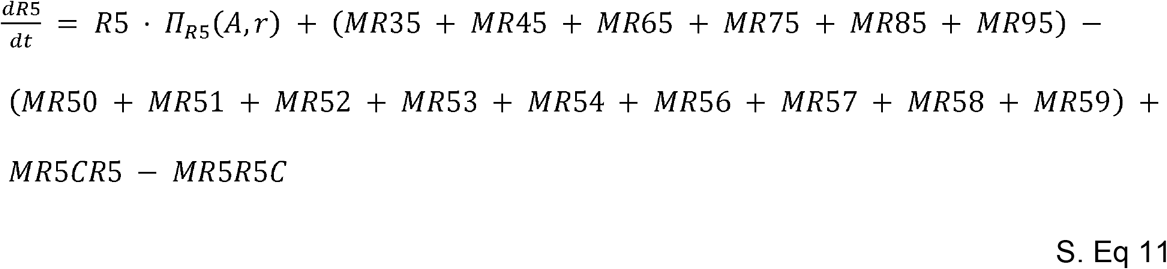

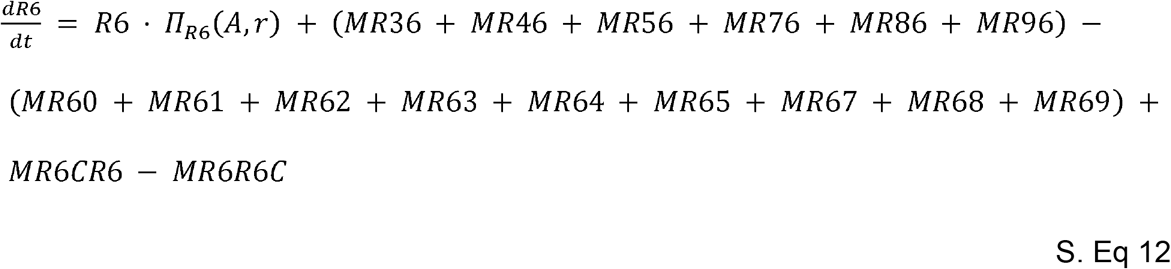

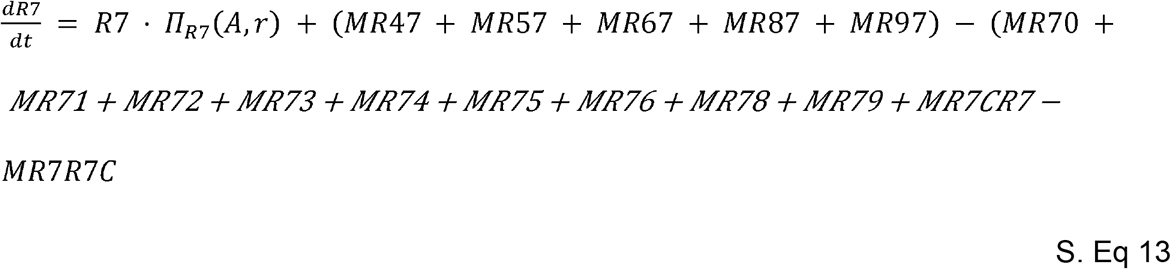

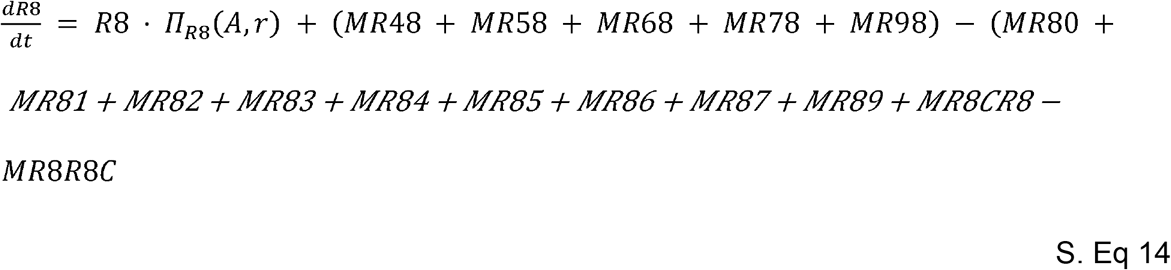

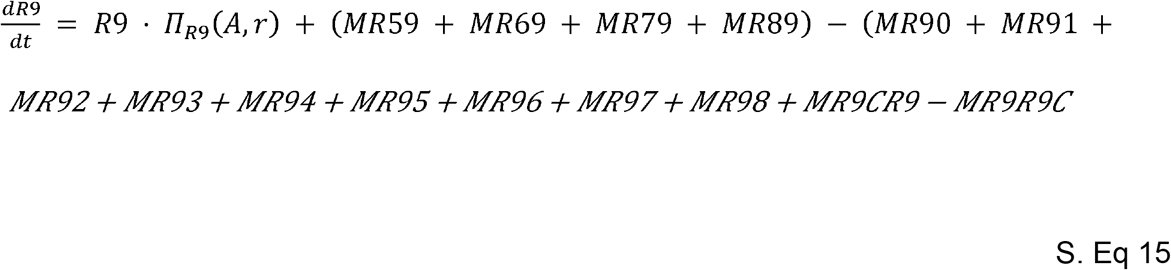

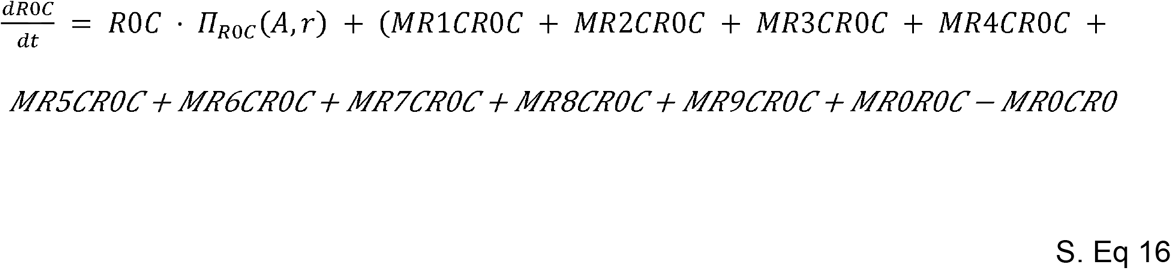

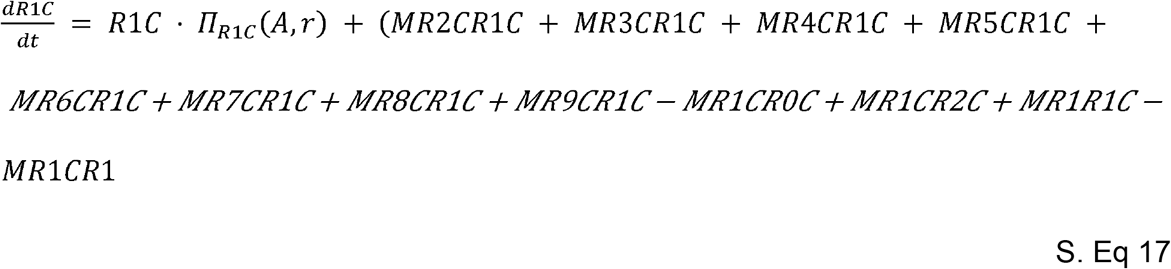

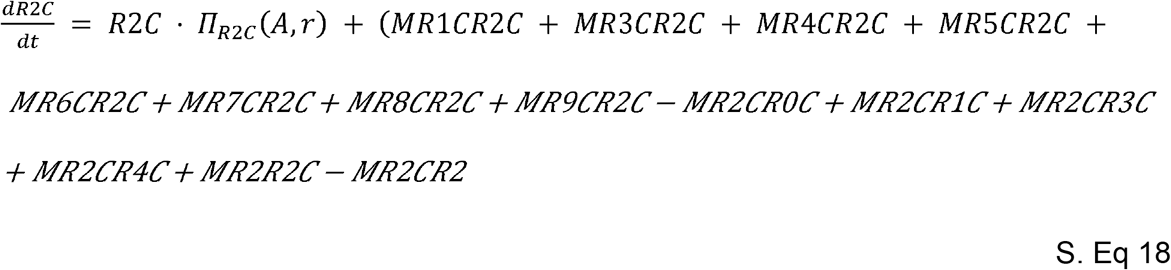

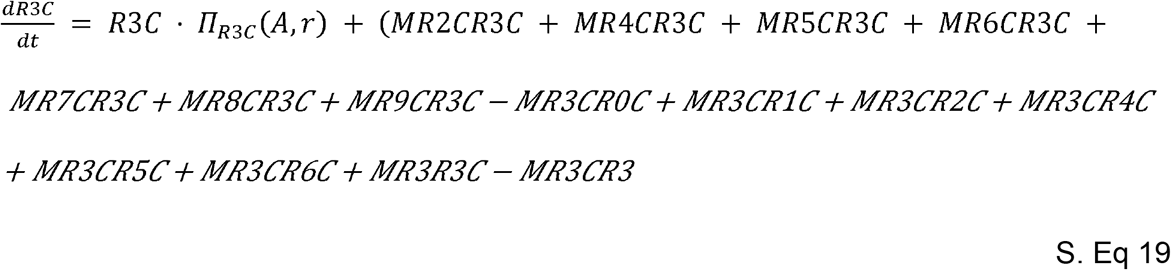

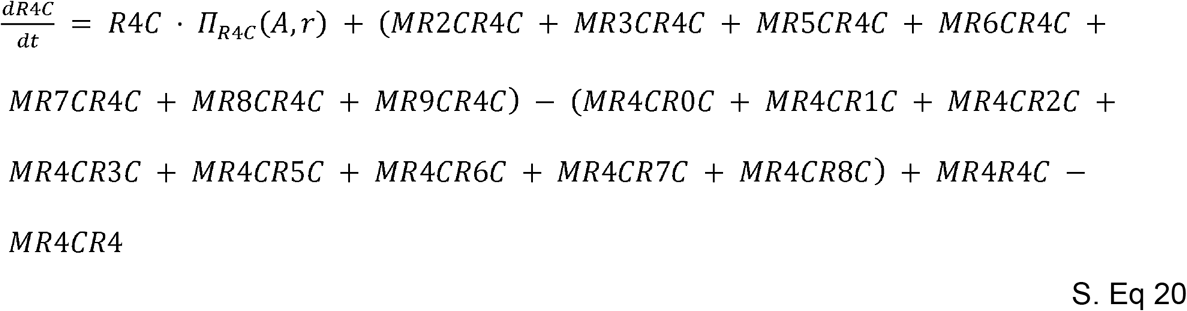

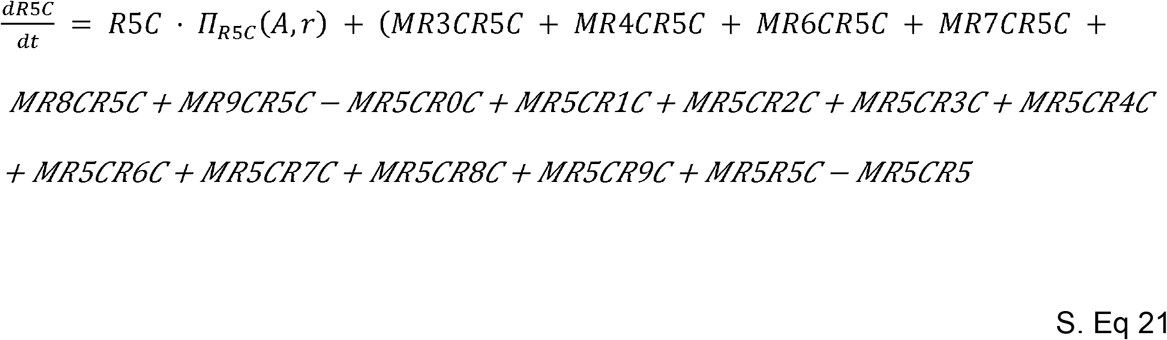

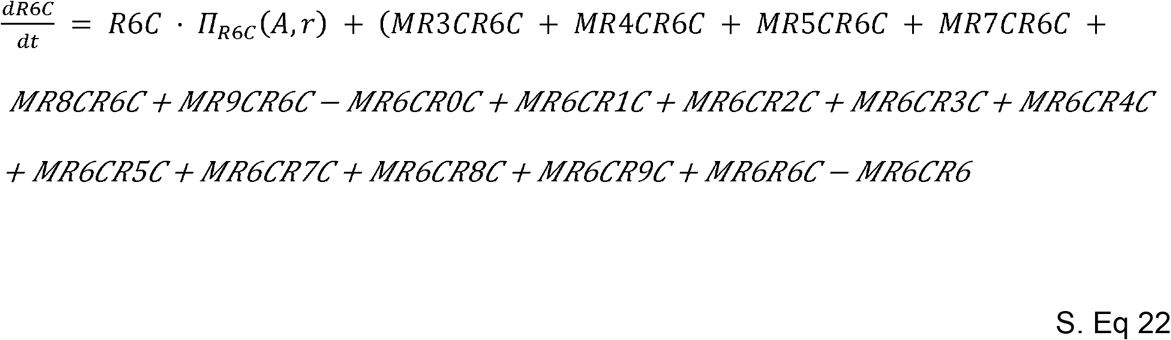

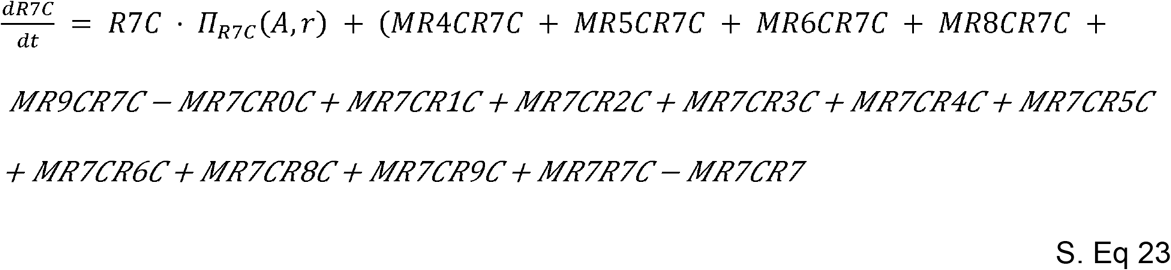

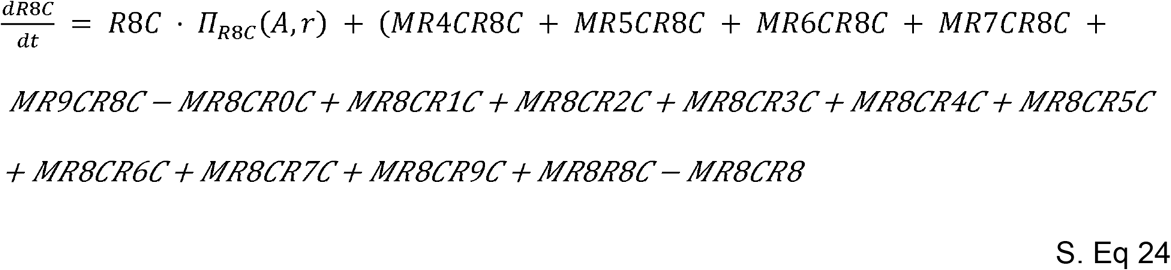

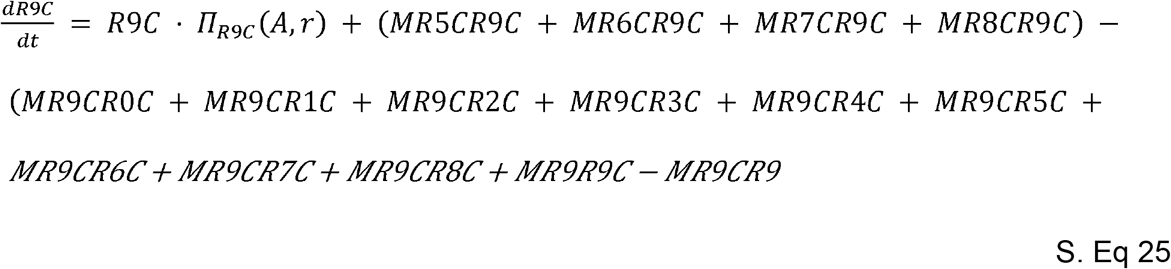

## Supplementary Figures

**Supplementary Figure 1.**
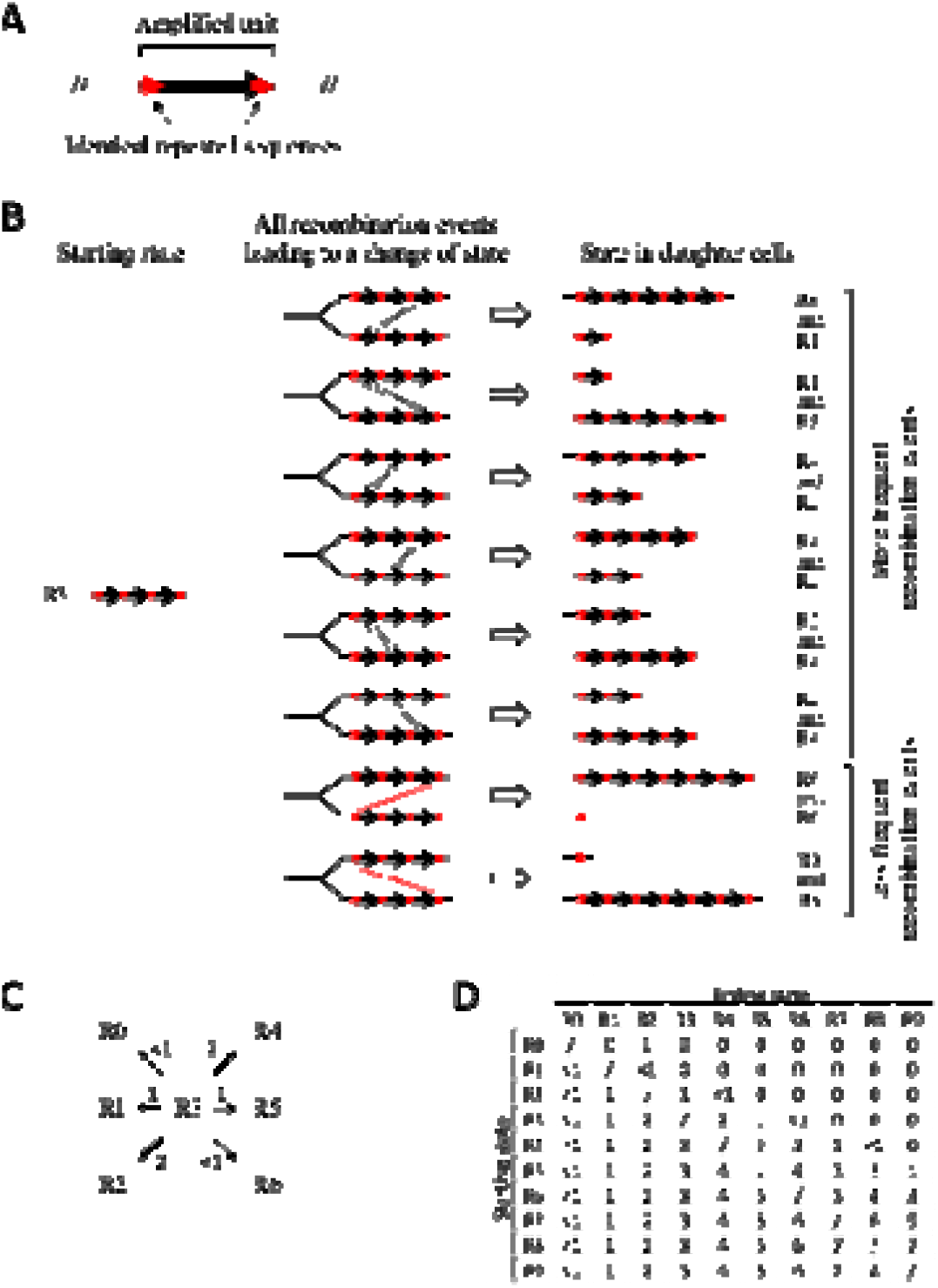
Transition and rates of transitions and mechanism within R states and RC states. Reproduced with permission from. ^10^**. A)** General structure of the amplified unit in our model. The amplified unit (often several kilobases in length) contains identical repeated sequences at each end (e.g., IS sequences <1 kb in length). **B)** Example, for cells starting with a triplication of the amplified unit (state R3), of all recombination events between amplified units present on sister chromatids and leading to a transition to other states in daughter cells. The more frequent recombinations occur between the amplified units (in black) while less frequent recombinations involve shorter identical repeats that are present at each end of the array of amplified units (in red). **C)** Summary of results presented in panel B schematizing the relative rates of transition between R states (arbitrary unit) for a cell in a starting state R3. **D)** All relative rates of transition between R states (arbitrary unit) for our model, for starting and ending R states constrained between R0 and R9.

**Supplementary Figure 2.**
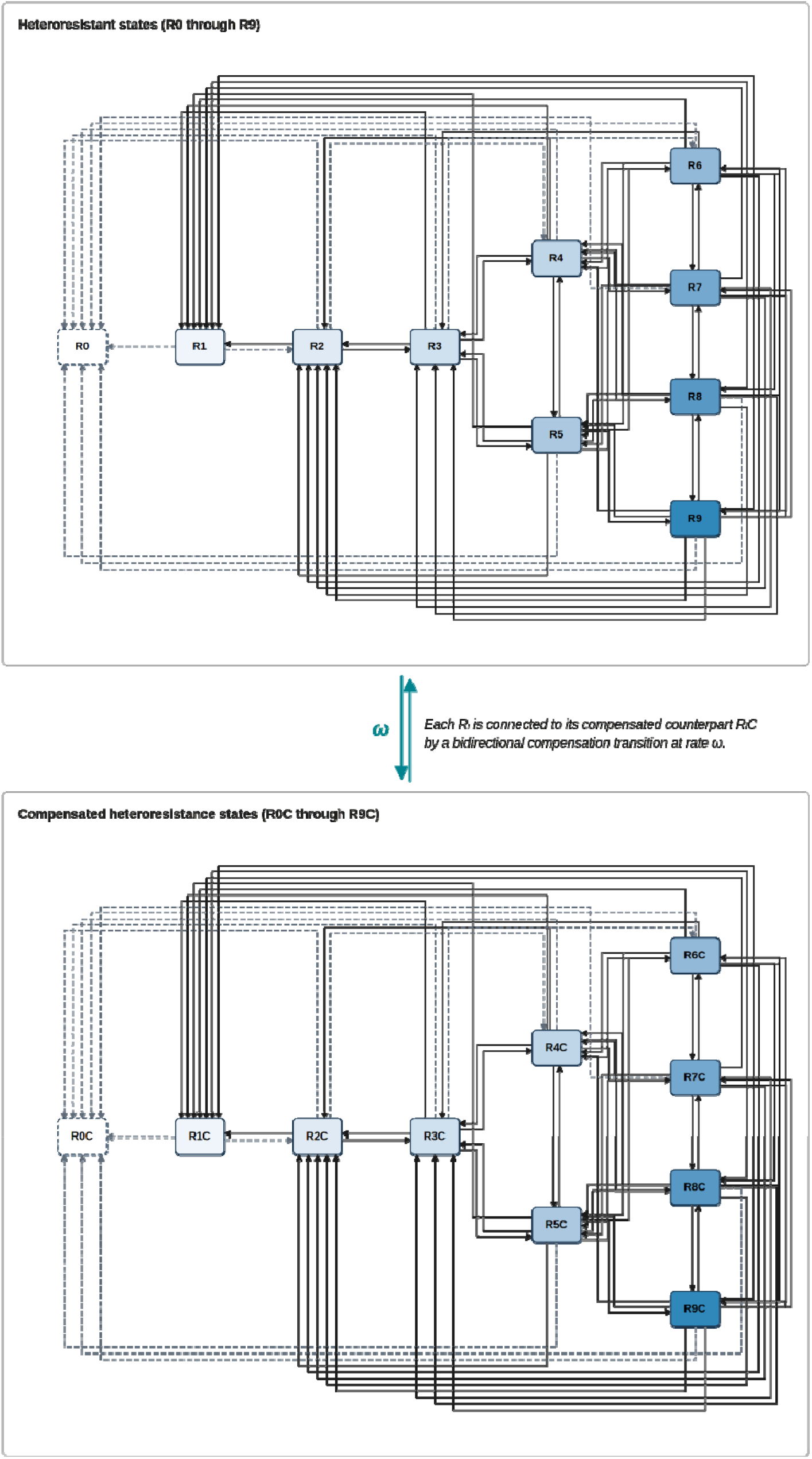
Diagram of the Model. The model comprises twenty bacterial states. R states (R0 to R9) correspond to bacteria with increasing copy number of a tandem amplified resistance gene, ranging from 0 to 9 copies; the compensated states (R0C to R9C) carry the same copy number but have additionally acquired a compensatory mutation. Solid arrows denote the more frequent recombination events (γ); dashed arrows denote the less frequent events (μ). Recombination occurs identically within the R space and within the C space. Teal arrows denote the compensation transition (ω) where each R state acquires a compensatory mutation, transitioning to its compensated counterpart. The model includes all possible transitions between states for starting states R0 to R9. The relative rates of all transitions are given in Supplementary Figure 1 and Supplementary Table 2.

**Supplementary Figure 3.**
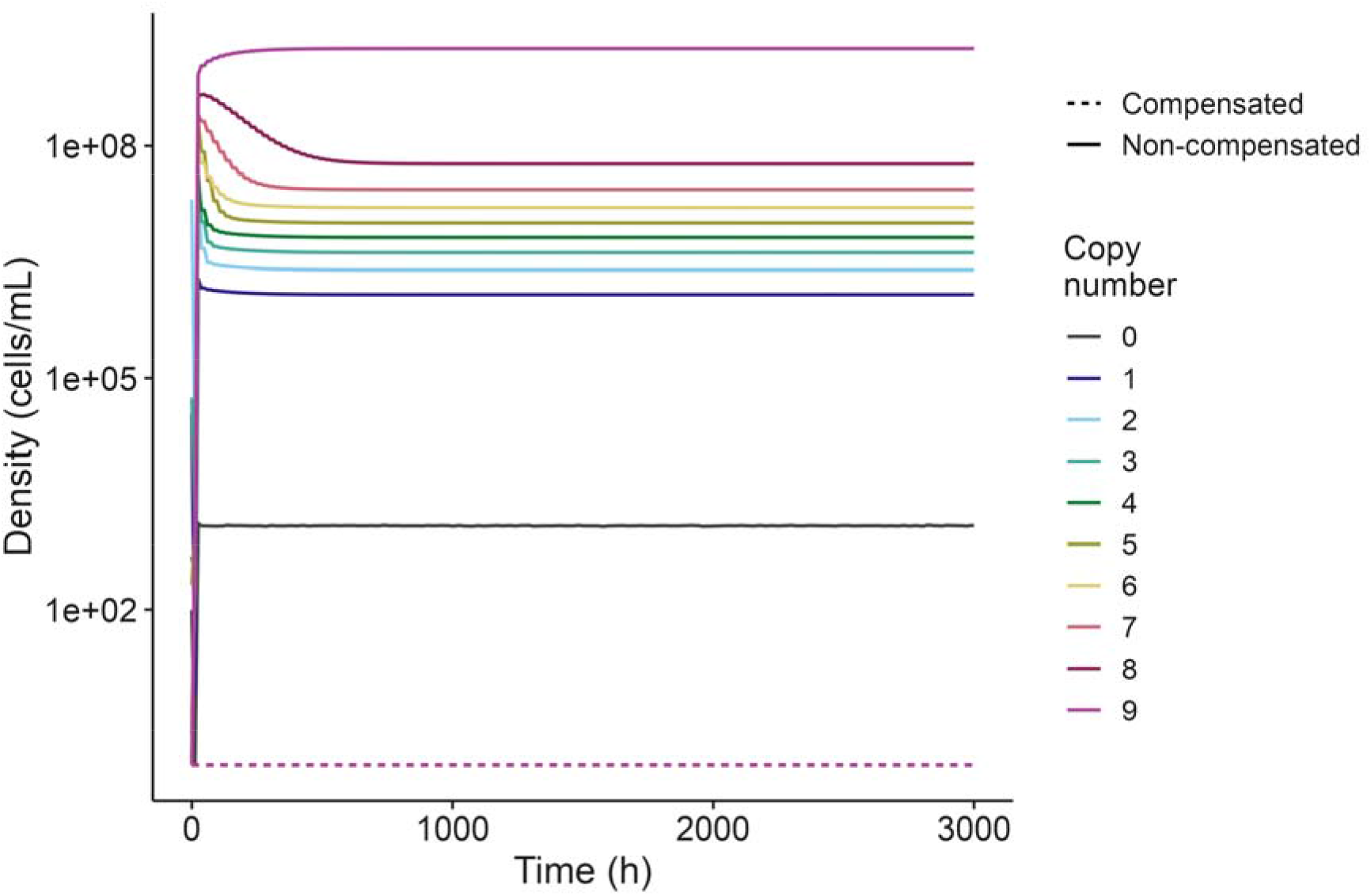
Dynamics of the model in the absence of compensatory mutations. The model was run under the conditions of Figure 1 with the compensation rate set to zero (ω = 0). The densities of the non-compensated states (R0 through R9) and the compensated states (R0C through R9C) are shown across the full 3,000 h simulation. The compensated states remain at zero, while the non-compensated states evolve by recombination alone and recapitulate the base model ^10^. Lines are coloured by copy number, with non-compensated states drawn solid and compensated states dashed.

**Supplementary Figure 4.**
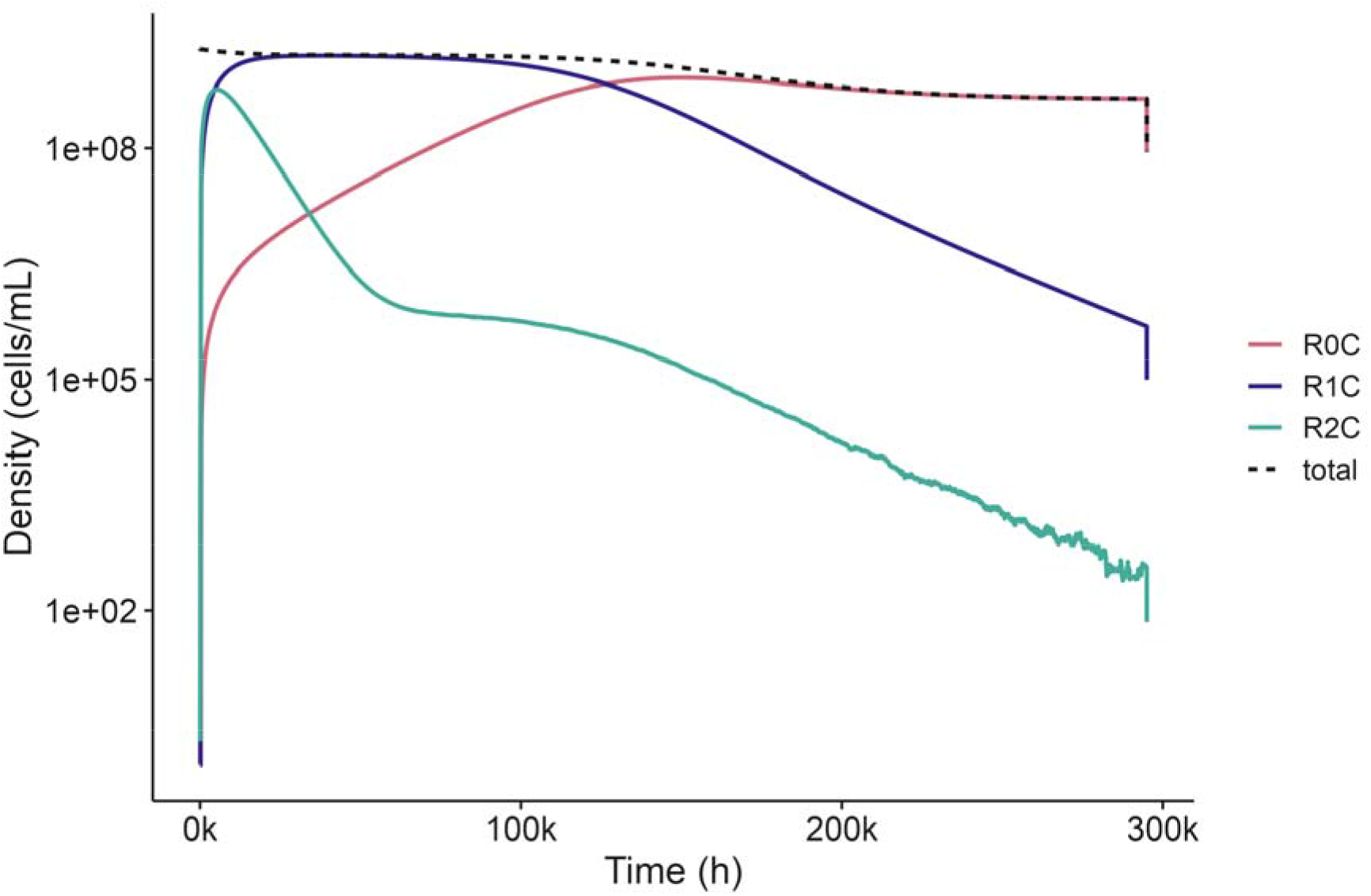
Far-horizon predictions over 300,000 hours. The deterministic model was run to 300,000 h to follow the final approach to the resistant endpoint. The three lowest compensated states, R0C, R1C, and R2C, and the total population (dashed black) are shown. The two-copy state R2C declines first, the single-copy state R1C dominates the intermediate horizon, and the zero-copy state R0C, the resistant endpoint that carries no amplification, overtakes R1C near 126,000 h—over 14 years, well beyond any clinically-relevant timescale.

**Supplementary Figure 5.**
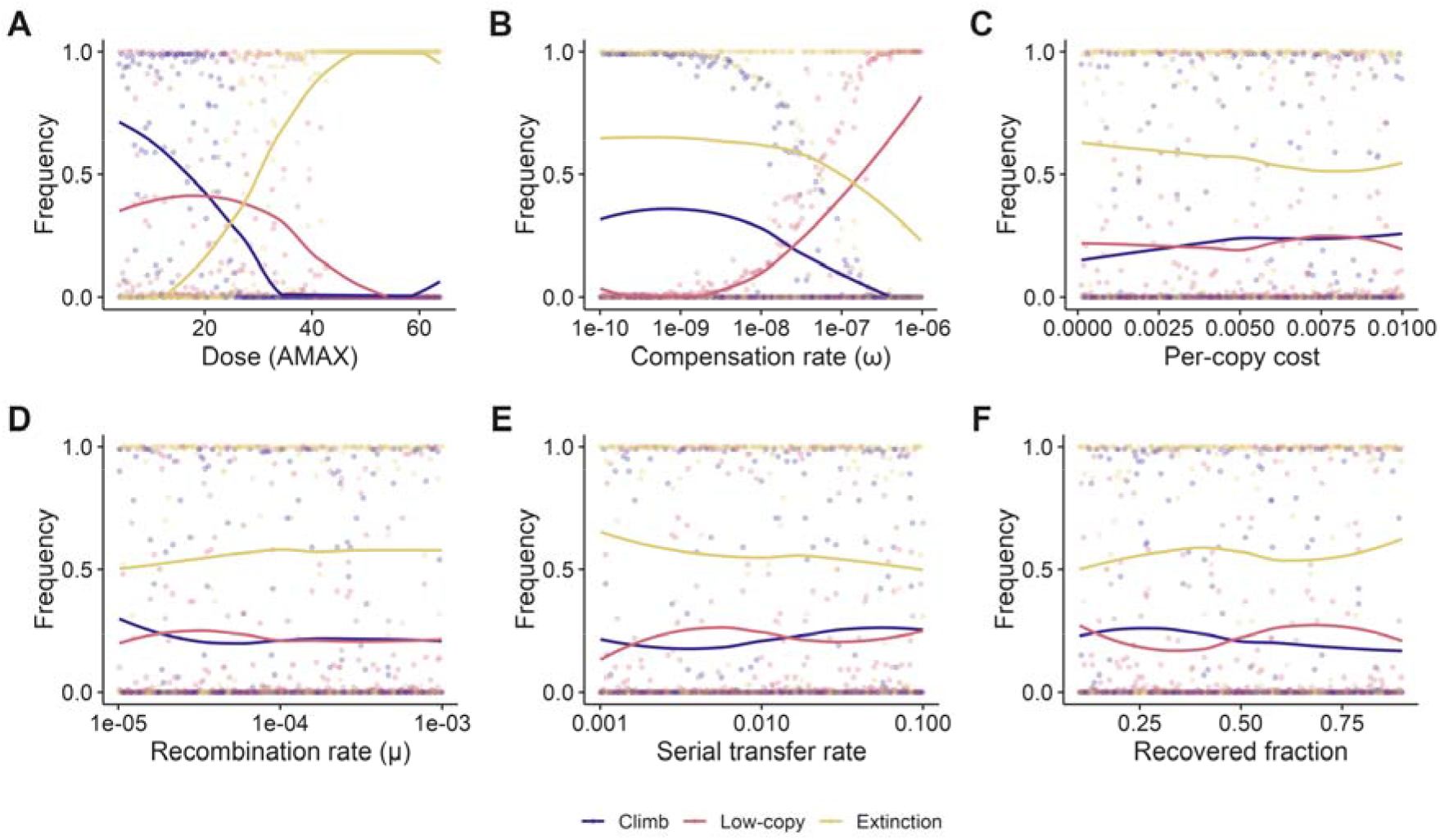
The raw parameter sweep. Each panel shows one of the six swept parameters on the x axis against the frequency of the three outcomes: climb, low-copy compensated survival, and extinction. Points are individual parameter sets and lines are smoothed trends. Dose drives extinction; the compensation rate governs the two survival routes; and the per-copy cost, recombination rate, serial transfer rate, and recovered fraction have little effect.

## Supplementary Tables

**Supplementary Table 1.** States in the Model.

| State | Dimensions | Definition |
| --- | --- | --- |
| r | µg per mL | Limiting Resource |
| R0 | Cells per mL | Antibiotic-susceptible Bacteria (Zero Copies of Resistance Gene) |
| R1, R2, R3, R4, R5,<br>R6, R7, R8, R9 | Cells per mL | Non-compensated Antibiotic-resistant Bacteria with Increasing Copies of Resistance Gene |
| R0C | Cells per mL | Compensated Antibiotic-susceptible Bacteria (Zero Copies of Resistance Gene) |
| R1C, R2C, R3C, R4C,<br>R5C, R6C, R7C, R8C,<br>R9C | Cells per mL | Compensated Antibiotic-resistant Bacteria with Increasing Copies of Resistance Gene |
| A | µg per mL | Antibiotic |
| B | Units per mL | β-lactamase |

**Supplementary Table 2.** Parameters in the Model.

| Parameter | Dimensions | Definition | Values | Source |
| --- | --- | --- | --- | --- |
| vR0,<br>vR1,<br>vR2,<br>vR3,<br>vR4,<br>vR5,<br>vR6,<br>vR7,<br>vR8,<br>vR9 | Per Cell Per<br>Hour | Maximum<br>Growth Rates | 2, 1.9987,<br>1.9974, 1.9961,<br>1.9948, 1.9935,<br>1.9922, 1.9909,<br>1.9896, 1.9883 | <sup>10</sup> |
| vR0C, vR1C,<br>vR2C, vR3C,<br>vR4C, vR5C,<br>vR6C, vR7C,<br>vR8C, vR9C | Per Cell Per<br>Hour | Maximum<br>Growth Rates of<br>Compensated<br>States | 2, 1.99935,<br>1.9987,<br>1.99805,<br>1.9974,<br>1.99675,<br>1.9961,<br>1.99545,<br>1.9948,<br>1.99415 | This<br>Study |
| $\mu_{Rx}$, $\gamma_{Rx}$<br>(frequent and rarer transitions) | Per Cell Per Hour | Recombination Transition Rates | $4 \cdot 10^{-4}$ to $4 \cdot 10^{-7}$ | <sup>10</sup> |
| $\omega$ | Per Cell Per Hour | Compensation Rate | $1 \cdot 10^{-7}$ | This Study |
| $v_{MIN}$ | Per Cell Per Hour | Minimum Growth/Death Rate | -2 | <sup>41</sup> |
| $\kappa$ | Dimensionless | Hill Coefficient | 1 | <sup>37</sup> |
| MICR0, MICR1, MICR2, MICR3, MICR4, MICR5, MICR6, MICR7, MICR8, MICR9 | $\mu\text{g per mL}$ | MIC of Each State | 0.375, 1.875, 3.375, 4.875, 6.375, 7.875, 9.375, 10.875, 12.375, 13.875 | <sup>10</sup> |
| MICR0C, MICR1C, MICR2C, MICR3C, | $\mu\text{g per mL}$ | MIC of Each Compensated State, Scenario 1 | 25, 25, 25, 25, 25, | This Study |
| MICR4C,<br>MICR5C,<br>MICR6C,<br>MICR7C,<br>MICR8C, MICR9C |  |  | 25,<br>25,<br>25,<br>25,<br>25 |  |
| MICR0C,<br>MICR1C,<br>MICR2C,<br>MICR3C,<br>MICR4C,<br>MICR5C,<br>MICR6C,<br>MICR7C,<br>MICR8C, MICR9C | µg per mL | MIC of Each<br>Compensated<br>State, Scenario 2 | 1.125,<br>5.625,<br>10.125,<br>14.625,<br>19.125,<br>23.625,<br>28.125,<br>32.625,<br>37.125,<br>41.625 | This<br>Study |
| pR0,<br>pR1,<br>pR2,<br>pR3,<br>pR4,<br>pR5, | Units per Hour | β-lactamase<br>Production Rate | 0.000, 0.001,<br>0.002, 0.003,<br>0.004, 0.005,<br>0.006, 0.007,<br>0.008, 0.009 | <sup>10</sup> |
| pR6,<br>pR7,<br>pR8,<br>pR9 |  |  |  |  |
| pR0C, pR1C,<br>pR2C, pR3C,<br>pR4C, pR5C,<br>pR6C, pR7C,<br>pR8C, pR9C | Units per Hour | β-lactamase<br>Production Rate<br>of Compensated<br>States | 0.000, 0.001,<br>0.002, 0.003,<br>0.004, 0.005,<br>0.006, 0.007,<br>0.008, 0.009 | This<br>Study |
| KB | Dimensionless | β-lactamase<br>Association<br>Constant | 1 | <sup>10</sup> |
| k | Dimensionless | Monod Constant | 1 | <sup>42</sup> |
| e | $\mu\text{g}$ | Conversion<br>Efficiency | $5 \cdot 10^{-7}$ | <sup>42</sup> |
| C | $\mu\text{g per mL}$ | Maximum<br>Resource | 1000 | Assumed |
| d | $\mu\text{g per Hour}$ | Antibiotic Decay | 0 | Assumed |
|  |  | Rate |  |  |
| AMAX | µg per mL | Antibiotic Dose at<br>Each Transfer | 16 | This<br>Study |
| dil | Dimensionless | Serial Transfer<br>Rate | 0.01 | This<br>Study |

